# TREND-DB – A Transcriptome-wide Atlas of the Dynamic Landscape of Alternative Polyadenylation

**DOI:** 10.1101/2020.08.04.235804

**Authors:** Federico Marini, Denise Scherzinger, Sven Danckwardt

## Abstract

Alternative polyadenylation (APA) profoundly expands the transcriptome complexity. Perturbations of APA can disrupt biological processes, ultimately resulting in devastating disorders. A major challenge in identifying mechanisms and consequences of APA (and its perturbations) lies in the complexity of RNA 3’end processing, involving poorly conserved RNA motifs and multi-component complexes consisting of far more than 50 proteins. This is further complicated in that RNA 3’end maturation is closely linked to transcription, RNA processing, and even epigenetic (histone/DNA/RNA) modifications. Here we present TREND-DB (http://shiny.imbei.uni-mainz.de:3838/trend-db), a resource cataloging the dynamic landscape of APA after depletion of *>*170 proteins involved in various facets of transcriptional, co- and posttranscriptional gene regulation, epigenetic modifications, and further processes. TREND-DB visualizes the dynamics of transcriptome 3’end diversification (TREND) in a highly interactive manner; it provides a global APA network map and allows interrogating genes affected by specific APA-regulators, and vice versa. It also permits condition-specific functional enrichment analyses of APA-affected genes, which suggest wide biological and clinical relevance across all RNAi conditions. The implementation of the UCSC Genome Browser provides additional customizable layers of gene regulation accounting for individual transcript isoforms (e.g. epigenetics, miRNA binding sites, RNA-binding proteins). TREND-DB thereby fosters disentangling the role of APA for various biological programs, including potential disease mechanisms, and helps to identify their diagnostic and therapeutic potential.

## INTRODUCTION

Next-generation RNA sequencing (RNA-seq) has led to the discovery of a perplexingly complex metazoan transcriptome architecture that results from the alternative use of transcription start sites, exon and introns, and polyadenylation sites (1; 2; 3). The combinatorial use and incorporation of such elements into mature transcript isoforms significantly expands genomic information and is subject to dynamic spatial and temporal modulation during development and adaptation. Recently, diversification of the transcriptome at the 3’end by alternative polyadenylation (APA) evolved as an important and evolutionarily conserved layer of gene regulation (4; 5). It results in transcript isoforms that vary at the RNA 3’end, which can have profoundly different physiological effects, by encoding proteins with distinct functions or regulatory properties, or by affecting the mRNA fate through inclusion or exclusion of regulatory elements, such as sites recognized by miRNAs or RNA-binding proteins (6).

Constitutive RNA 3’end processing relies on a complex macromolecular machinery that catalyzes endonucleolytic cleavage and polyadenylation (CPA) of pre-mRNA molecules (7). This involves the assembly of four multi-component protein complexes (CPSF, CSTF, CFI, and CFII) (8) on the pre-mRNA at dedicated, but largely poorly conserved, processing sites (9). Differential expression of individual complex components (10; 11; 12) or regulation of their binding properties (e.g. by signalling (13)) can direct the dynamic modulation of APA resulting in transcript isoforms with alternative 3’ends. In addition, other mechanisms including RNA polymerase II kinetics (14; 15) or epigenetic events (16; 17; 18; 19; 20; 21; 22) regulate APA, illustrating a previously unanticipated complex crosstalk between various cellular processes in the control of transcriptome diversity. Although difficult to detect by standard high throughput profiling techniques (23; 24; 25), dynamic changes at the transcriptome 3’end are widespread (26; 27; 28; 29). They affect more than 70% of all genes. Global APA changes are commonly associated with differentiation and de-differentiation processes (30). This requires a well-coordinated temporal and spatial interaction between RNA motifs and the multi-component processing complex to ensure that CPA occurs timely and at the right position (31; 32). Hence 3’end processing is tightly coupled to transcription and other co- and posttranscriptional processes (7; 33), and controlled by delicate (auto-) regulatory mechanisms (34; 35). Transcripts that show dynamic regulation at the 3’end are typically encoded by phylogenetically old genes, which corresponds to the phylogenetic age of most executing APA regulators (12). Finally, individual components of the CPA machinery are epistatic during differentiation and de-differentiation; they display functional dominance over ‘neighboring’ factors within and across the multi-component CPA complexes in the control of APA (12).

While the underlying mechanisms of APA regulation are still being elucidated, disruption of this process clearly proved to be clinically relevant (36). For example, APA perturbations are associated with various disorders (37; 11; 38; 39; 40; 41) (and refs therein), but they can also possess direct disease eliciting activities, act as oncogenic driver, and thereby mimic genetic alterations (12; 42). Importantly, such changes are generally missed in genome profiling endeavours (43). But often they also remain undetected by standard RNA-seq technologies (44). However, even when resulting in primarily subtle changes of non-coding RNA sequence elements in the 3’UTR, APA perturbations can be functionally significant (12) and represent unexpectedly potent novel biomarkers (24; 12). Aberrant posttranscriptional expansion of the genome complexity can thus result in most devastating consequences; at the same time, it also opens novel diagnostic (24; 12) and therapeutic avenues (45; 46).

In an attempt to systematically identify drivers of APA in tumorigenic differentiation/de-differentiation processes (12), we recently applied a comprehensive RNAi screening coupled to a high-throughput sequencing approach suited to capture polyadenylated transcript 3’ends of numerous experimental conditions (TRENDseq) (44). We depleted more than 170 proteins, including all known factors involved in pre-mRNA 3’end CPA in eukaryotes (8) and selected key factors regulating transcriptional activities, splicing, RNA turnover, and other functions (47; 48; 49; 50; 51; 52; 53; 54; 55; 56; 57; 58; 59) which could directly or indirectly affecting APA (5; 60). This resulted in the identification of numerous drivers directing APA including the transcription termination factor PCF11 with a critical role in neurodevelopment and tumor formation (12).

Here, we present TREND-DB, the full roll-out of a user-friendly database, which in complementation to existing static polyA DBs (61; 62; 63; 64) (and refs therein), catalogs the dynamic nature of the APA landscape genome-wide upon targeting *>*170 components involved in the definition of RNA 3’ends. We provide a highly interactive APA network map displaying central hubs in APA regulation. TREND-DB allows querying effects of selected APA-regulators including condition-specific Gene Ontology (GO) (65) enrichment analyses of APA-affected genes. Including an instance of the UCSC Track Hub framework also allows the investigation of other gene regulatory mechanisms with reference to the transcriptome-wide APA data (e.g. miRNA and RNA-binding proteins, epigenetics), or alternatively the direct comparison with own data, uploaded to the genome browser. TREND-DB thereby helps to illuminate the role of APA for various biological programs and potential disease mechanisms, and allows to identify their diagnostic and therapeutic potential. The functional enrichment across all knockdown conditions already suggests wide biological and clinical relevance.

## DATA COLLECTION AND PROCESSING

### TRENDseq data collection and processing

Applying TRENDseq combined with an RNAi screening targeting *>*170 potential APA regulators (12), we previously identified more than 9.000 APA-events (out of approximately 20.000 expressed transcripts), corresponding to more than 3.600 genes significantly affected by APA. Briefly, to this end we first generated a TREND annotation assembly based on 3’end reads to identify bona fide polyadenylation events (23). This annotation was used to exclude TRENDseq reads originating from internal priming on genome encoded adenosine-rich regions. TRENDseq data originating from the RNAi screening were demultiplexed, trimmed, mapped to the human hg38 genome, and filtered from internal priming events using the TREND annotation (for further details, see (12) and Supplementary Methods).

### Quantification and global characteristics of APA events

The number of reads aligned to each polyadenylation site is used as a proxy for the expression of an individual 3’end transcript isoform. The expression level of each transcript isoform was examined by Fisher’s exact test in comparison to the respective other 3’end isoform(s) expressed by the same gene. A contingency table included the number of reads of the tested isoform and total amount of reads of all the other isoforms of the gene (for the knockdown and control samples, respectively). Obtained p-values were adjusted using the Benjamini-Hochberg method, and a threshold on the adjusted p-value *≤*0.05 filter was applied (for further details on normalization, fold change of regulation, and calculation of the shortening index see (12)).

Interestingly, almost all depleted factors showed APA-regulation on average affecting 130 genes (Fig. 1) with a mean distance between APA-regulated sites of approximately 1.020 nucleotides (Supplementary Table 1). While the vast majority of APA events (65%) occur in the 3’UTR (‘tandem polyadenylation’), 20% of the events affect internal polyadenylation (i.e. within introns or the coding region), thereby modulating the generation of truncated protein isoforms. Directly comparing the effects quantitatively and qualitatively, we observed that APA is predominantly driven by components of the CPA machinery (38% of all APA events, with key factors affecting up to 1.400 genes (e.g. NUDT21, CPSF6, and PCF11), average effect 319 genes) and that their function is mainly non-redundant (12). However, a significant proportion of APA (62%) is controlled by components involved in transcription (14%, on average affecting APA of 133 genes) and other co- and post-transcriptional events (e.g. splicing, RNA-turnover (16% and 9%; on average affecting APA of 128 and 89 genes, respectively)) or epigenetic modification. This corresponds to a quantitatively comparable extent to which individual splicing factors influence alternative splicing (66). Notably, while the depletion of the vast majority of factors results in even amounts of shortened and lengthened transcript isoforms, the depletion of the CFI and CFII complex components (CPSF6, NUDT21, and PCF11) directs APA in an almost exclusive unidirectional manner resulting in uniformly shortened (CPSF6 and NUDT21) or lengthened (PCF11) transcript isoforms (Fig. 1B). In particular, our screening also identified APA regulation to be associated with factors involved in genome surveillance or known to drive tumor-suppressive programs (e.g. TP53), as well as other processes involved in the coupling between oncogenic signals and 3’end processing, such as BARD1 (67).

**Figure 1:**
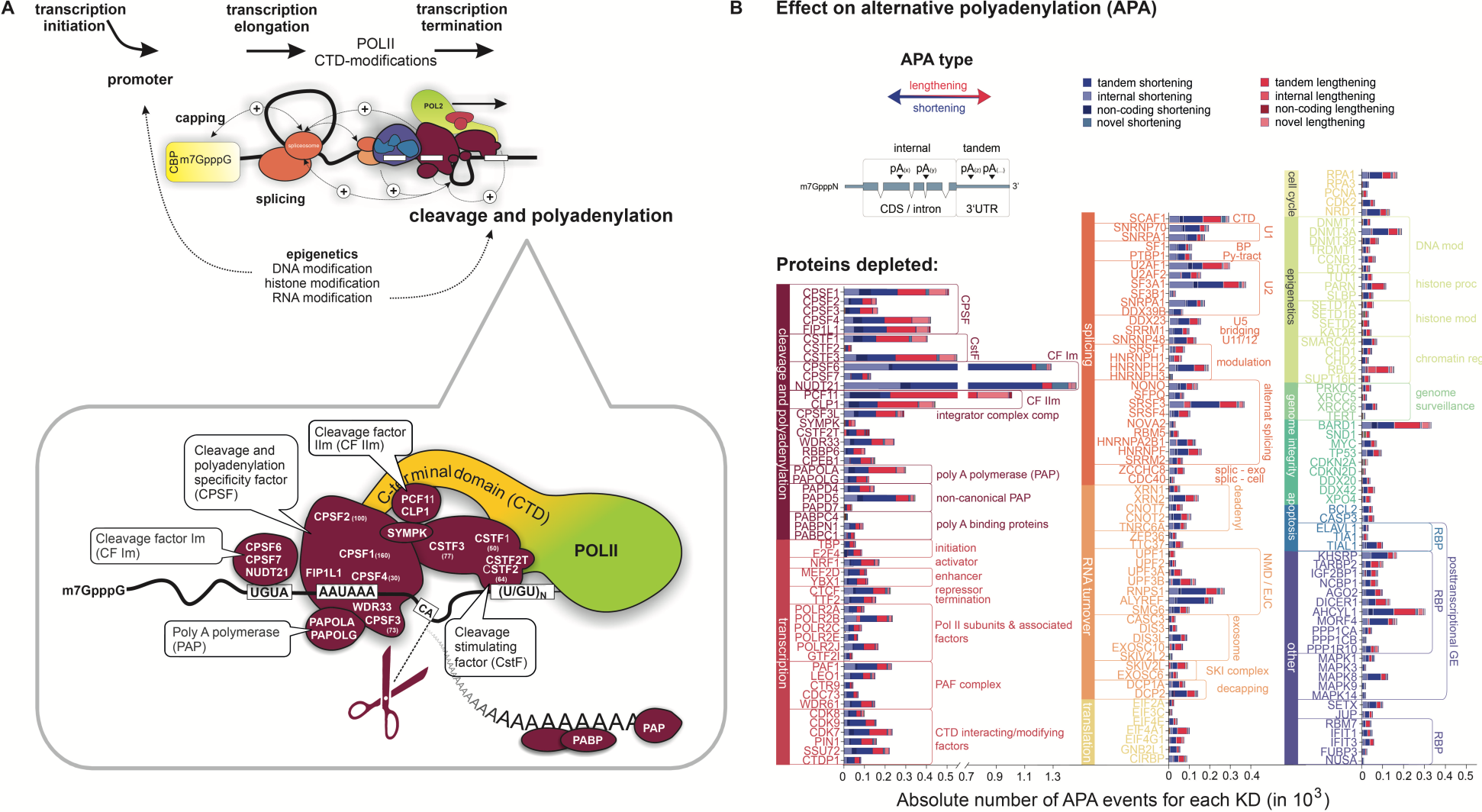
Data preview of TREND-DB cataloging the dynamic landscape of alternative polyadenylation **(A)** 3’end processing of polyadenylated RNAs relies on a complex macro-molecular machinery that catalyzes endonucleolytic cleavage and polyadenylation (CPA) of the nascent pre-mRNA molecule. CPA is carried out after assembly of four multi-component protein complexes (CPSF, CSTF, CFI, and CFII) interacting with multiple RNA motifs. This is tightly coupled to transcription, other steps of RNA processing, and even epigenetic modifications. Differential expression of individual complex components or regulation of their binding properties can redirect CPA to alternative processing sites. In addition, further mechanisms including RNA polymerase II kinetics can regulate alternative polyadenylation. **(B)** Global overview of alternative polyadenylation (obtained by transcriptome-wide TRENDseq) after depletion *>*170 components involved in various facets of transcriptional, co- and posttranscriptional gene regulation, epigenetic modifications, and further processes with a potential role in the definition of RNA 3’ends. APA shortening and lengthening refers to CPA at an upstream or downstream positioned poly(A) site (PAS). This can modulate CPA at alternative PASs located in the 3’UTR (“tandem” APA) or in the coding sequence (CDS) and/or intronic region (“internal” APA), resulting in transcript isoforms with distinct regulatory properties or encoding c-terminally modified proteins. TREND-DB is designed to provide interactive and accessible exploration of this data set.

## DATABASE CONTENT AND USAGE

To make the dynamics of APA accessible to a wider scientific community, we constructed a user friendly, highly interactive database. In the database, we store (1) Summary files for each of the 175 knockdown experiments, where effect size (reported as fold changes) and significance (as adjusted p-values) of each isoform, together with the shortening index (SI), which describes the overall tendency of a given gene to express shortened (or lengthened, respectively) transcript isoforms (based on a proxy of the two most significantly affected APA isoforms), are reported; (2) bigWig files with the coverage for the isoform peaks, used to compute the SI values across conditions; (3) BED files with genomic positions for curated lists of poly(A) sites and known microRNA binding sites for the hg38 assembly (Fig. 2, upper section).

**Figure 2:**
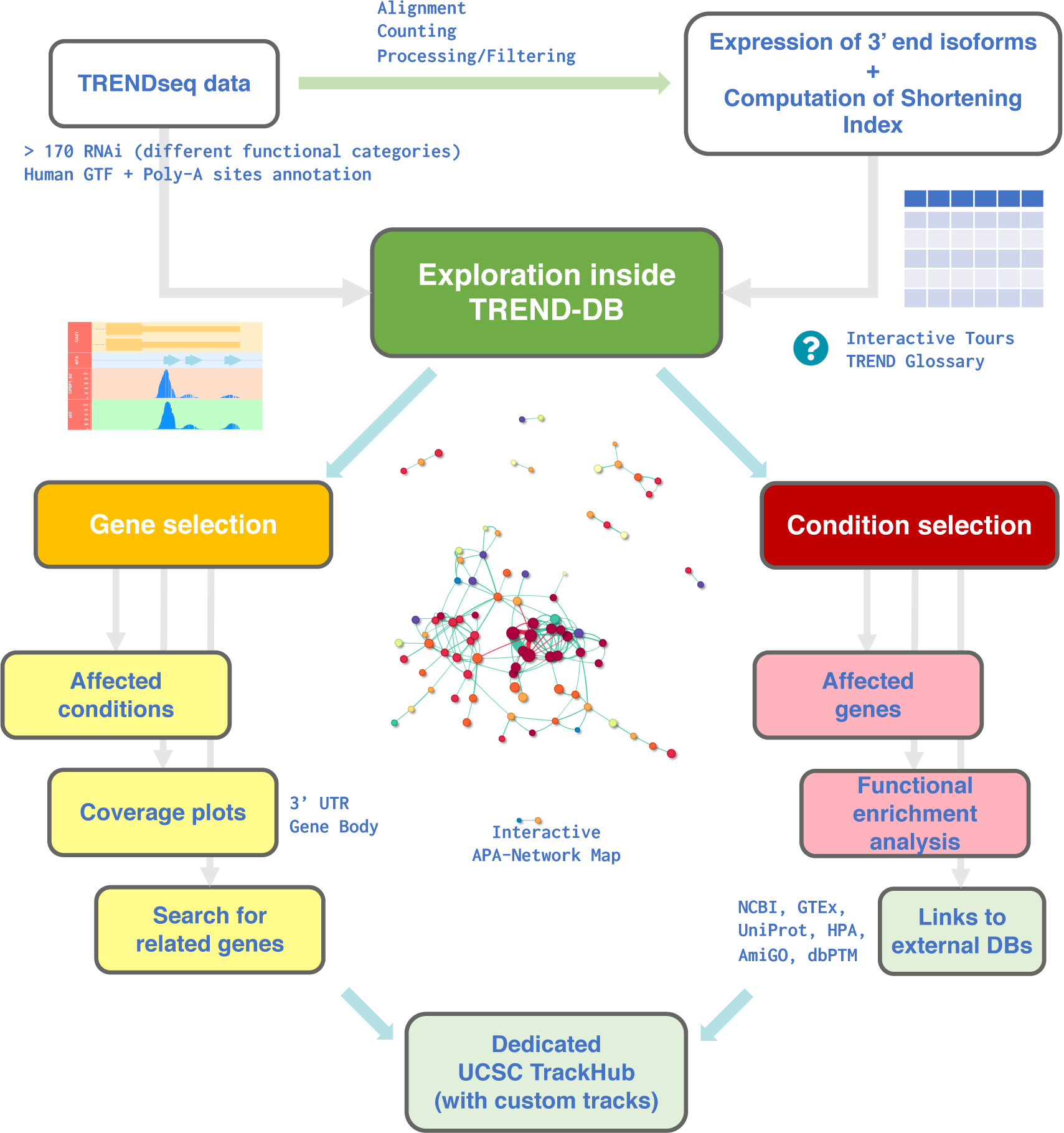
Overview of the exploration workflow with TREND-DB. TRENDseq data were aligned, quantified, and processed (12) to obtain the expression values of APA transcript isoforms, and the corresponding Shortening Index (SI) across all siRNA-conditions (displayed in the ***Data Preview*** tab). The ***Main View***, where an interactive APA-network map is displayed, enables users to inspect both dimensions of the underlying resource. Upon selection of a gene (left, ***Gene View***), all siRNA conditions affecting APA of this gene are displayed, accompanied by the TRENDseq coverage plot centered on the desired region (3’UTR or gene body); genes showing a similar APA phenotype (based on the SI profiles) can also be retrieved. When selecting a siRNA-condition (right, ***Condition View***), all APA-affected genes are listed, and users can perform functional enrichment analysis on the fly. External databases can be reached via dynamically generated action buttons, reducing the time to retrieve additional information. A dedicated instance of the UCSC Browser is provided, containing all the genomic tracks of the TRENDseq dataset, with the possibility to integrate custom tracks.

The definition of the software requirements has been established to cover the typical use cases, where the users can view, filter, and visualize the underlying data. An overview is followed by dedicated gene-centered and condition-centered views, offering the possibility to store the generated visual and tabular outputs (Fig. 2).

The TREND-DB application is developed in R (68) as a Shiny web application (69), leveraging the reactive programming paradigm offered by this framework. This framework has been chosen over existing other ecosystems such as BioPyramid (70) for its simplicity and efficiency of reactivity, together with the customizability of the analysis components enabled by the R programming language. Moreover, we leverage on a number of core packages and infrastructures, such as GenomicRanges (71), from the Bioconductor framework, which efficiently performs a variety of operations in omics data contexts (72). Tabular information is displayed in TREND-DB as interactive tables, generated with the DT package (73), as an R-based interface to the JavaScript DataTables library, which contains useful features, such as pagination, sorting, and filtering. Plots of genomic regions (3’UTR and gene bodies) for selected features are implemented with the Gviz package (74).

### The user interface

The user interface is based on a dashboard page structure from the shinydashboard package (75), where the main functionality is divided into a series of tab-based views (represented by the blocks in Fig. 2).

The ***Data Preview*** tab serves as a starting point for exploring the provided data set (see also the dedicated guided tours in the TREND-DB web browser). It offers an overview table that contains all genes and shows their shortening index (SI) values in all involved knockdown conditions, enabling to filter for defined SI thresholds applied on absolute values. Selecting a gene in the overview table prints an additional table, listing all the knockdown conditions in which it is affected.

The ***Main View*** tab first displays an interactive view of the APA network, implemented with the visNetwork package (76). The APA network map (Fig. 2 and Fig. 3A) essentially depicts cooperative (and antagonistic) interactions between APA regulators (affecting APA of identical genes). While the radius of the nodes reflects the number of APA affected genes (per condition), the links between a pair of nodes depict synergism (green, i.e. unidirectional lengthening or shortening) or antagonism (red, i.e. reciprocal lengthening or shortening) on APA of the common pool of genes, with the edge width encoding the significance of the overrepresentation of genes being synergistically or antagonistically regulated (BH-adjusted p-values)(12). Remarkably, the network layout reflects known protein complexes involved in polyadenylation (8) and the crosstalk to other RNA processing events (7), recapitulating the underlying molecular architecture.

**Figure 3:**
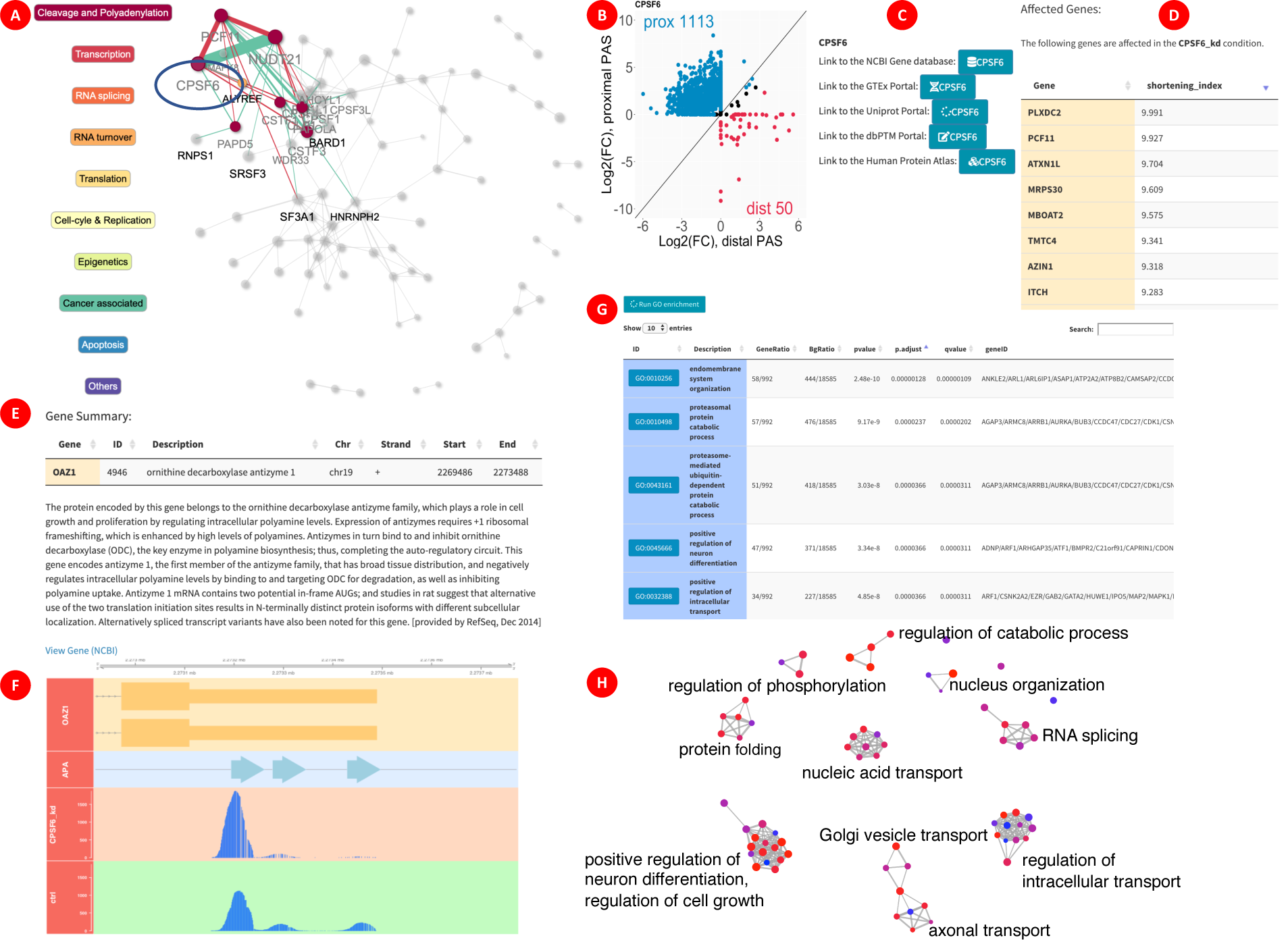
Selected screenshots to illustrate the exploration of the APA landscape with TREND-DB. From the ***Main View*** tab, users can select any condition among those included in the RNAi screening, either from a dropdown menu or from the interactive APA network **(A)**. For example, upon selection of CPSF6 (component of the CFIm protein complex, highlighted in the circle) ‘neighboring’ components/complexes that also regulate identical APA-affected target genes are highlighted. In this case CPSF6 was found to regulate APA with a global shortening bias, as displayed by the scatterplot in **(B)** summarizing the PAS usage. Links to external databases **(C)** are automatically displayed for simplifying further exploration steps beyond TREND-DB. Moreover, a table containing the affected genes (sorted by the Shortening Index in **(D)**) is displayed, and can be used to further explore the APA phenotype of a particular gene - e.g. OAZ1, both in the ***Main View* (E)**, where general information is shown, and in the ***Gene Plot*** tab, where users can inspect the coverage tracks for a locus of interest, such as the 3’UTR, where the effects of isoform shortening or lengthening is shown **(F)**. Using the whole set of affected genes in a particular screening condition (i.e. knockdown), TREND-DB enables the direct calculation of functional enrichment of APA-affected target genes. The functional enrichment is displayed in an interactive table **(G)**, or graphically as an enrichment map **(H)** for easier interpretation (for improving the readability, the single node labels have been replaced with a representative biological process for the clustered subset).

The remainder of the tab is separated into two sections, corresponding to the two dimensions of the database: (1) a ***Gene View*** and (2) a ***Condition View*** (Fig. 2, central section). Any gene or condition selection in the *Data Preview* tab is automatically carried over to this tab, while a (different) gene or condition can also be selected from the respective dropdown menu. Upon selecting a gene, the ***Gene View*** section displays a brief summary of the gene, including name, coordinates, strand, chromosome, and a description (Fig. 3E). Hyperlinks to additional resources, such as the NCBI RefSeq database, the GTEx Portal, the Human Protein Atlas, or the dbPTM database (77; 78; 79; 80), are provided as well to facilitate further exploration. On the other hand, once an experimental condition has been selected, the ***Condition View*** provides a list with all genes affected, their SIs, and p-values for the shorter and longer isoforms, according to the APA usage (Fig. 3D). Users may view a scatterplot for each condition, depicting the distribution of proximal and distal poly(A) site usage amongst the affected genes (Fig. 3B).

Additionally, a built-in ***GO term enrichment analysis*** tool (Fig. 3G) performs functional analysis on the set of genes affected in the selected condition against a background of all genes, as a potential starting point for more in-depth analysis - the interactive table created in TREND-DB automatically links to the corresponding entries in the AmiGO database (81), and can be represented visually as an enrichment map for better overview of the impacted biological themes (82). For example, genes affected in the AGO2 knockdown display among the top enriched results three retina-related GO terms, a tissue where 3’UTR shortening is often observed (83). However, there are also other interesting examples of GO enrichments underlining the wide biological and clinical relevance across knockdown conditions, for example suggesting either a potential bidirectional mechanistic link between alternative polyadenylation and alternative splicing, but also mechanisms in UTR biology towards immunobiology or developmental processes (Fig. 3H).

We and others earlier demonstrated that the relative abundance of transcript isoforms generated by APA has strong diagnostic (prognostic) potential, with superior performance in predicting death or high risk in cancer patients compared to commonly used clinical stratification markers (12). In addition, disentangling the functional role of (select) transcript isoforms opens ample therapeutic opportunities to interfere with polyadenylation in a selective or non-selective fashion (45; 46). This requires further in depth information on where APA occurs. We thus implemented functions that allow to visualize where APA affected genes are differentially polyadenylated, whether this affects the coding region of the gene (‘internal polyadenylation’) and/or non-coding UTR elements (‘tandem polyadenylation’, see above) and whether there are other conditions that result in a similar APA ‘pattern’ of the same gene.

The ***Gene Plot*** tab creates overview plots of a gene and its associated data (Fig. 3F). These plots contain a gene region reference track, coverage tracks for the selected condition (otherwise defaulting to the condition where the maximum SI is observed) as well as the control, and a genomic track for known APA sites. By default, overview plots are centered around the 3’UTR region of the gene; however, one can display the whole gene body region in the plot. Generated plots can be downloaded using a download button. Similarly affected genes, suggested in the interface component below the plot, can also be selected and displayed.

To enable a seamless integration with the wealth of genomic information provided by other sources, such as the UCSC Genome Browser (84), we set up an instance of the UCSC Track Hub (85), where the data is stored on a secure FTP server and the required configuration files (hub.txt, genomes.txt, and trackDb.txt) are specified in the TREND-DB GitHub repository, specifying the properties of the hub data tracks (genome assemblies, name, labels, descriptions, and display parameters). This content is integrated in the ***Genome Browser*** tab of TREND-DB via an iframe HTML element (for displaying web pages within other web pages). A custom URL is generated, accounting for all current selections made by the user, and passes it to the iframe, where the Genome Browser mirrors the behavior of the TREND-DB application.

While the ***Gene Plot*** view has a more specific aim on visualizing TRENDseq data (e.g. whether APA primarily affects the 3’UTR or the coding region), the ***Genome Browser*** view constitutes a bridge to a more flexible environment, where users can upload additional resources, such as existing tracks for their own genomic data, and facilitate the integration with the dataset of TREND-DB. As a proof-of-concept we implemented miRNA tracks; however, this can also be expanded to other resources e.g. CLIPseq (86), global interactome data (87; 88), or DNA/RNA modifications (19; 21; 22).

Altogether, these functions enable accessible and extendable exploration for the TREND-DB data sets, for unbiased and targeted analysis of downstream functions, for assessment of the diagnostic potential of APA signatures, and for the development and design of potential therapeutic strategies (89). TREND-DB thereby facilitates access and reuse of such complex data by a broader scientific community complying with the FAIR data principles (90).

### Simplifying the exploration for users

A common issue when exploring a series of data with a new application is how to efficiently guide the users across the different pieces of functionality offered. While text-based tutorials and walk-through documents (also in video format) can work efficiently for a specific snap-shot of the application, it is not a convenient approach for maintenance once new features are added. Moreover, an approach which encourages the users to try out specific actions tightly linked to defined elements in the user interface has proven useful for other applications such as ideal (91) and iSEE (92), offering a learning-by-doing paradigm for showcasing the application features in a more cohesive, workflow-like way.

Each tab of TREND-DB has a dedicated question mark action button, which triggers bespoke step-by-step tours of the functionality offered in each view, via the rintrojs package (93). We guide the audience in the original documentation by displaying use cases that refer to a few selected findings reported earlier (12).

Moreover, as TRENDseq data is not as common as e.g. transcriptome datasets, we included a Glossary section in the main dropdown menu, precisely defining the terms commonly used throughout the app, and specifically referring to the data type under inspection.

### Availability of database, code, and data

The TREND-DB application has been deployed on the Shiny Server at the Institute of Medical Biostatistics, Epidemiology and Informatics (IMBEI), and can be accessed at http://shiny.imbei.uni-mainz.de:3838/trend-db.

The data (sequencing raw and processed) displayed in TREND-DB is also available on GEO, under the accession GSE95057 https://www.ncbi.nlm.nih.gov/geo/query/acc.cgi?acc=GSE95057).

TREND-DB is freely available as a Shiny web application, with source code and processing information available under the MIT license at https://github.com/federicomarini/trend-db.

## SUMMARY AND FUTURE DIRECTIONS

TREND-DB has been conceived as an application to enable easy access to the main datasets from a recently published massive RNAi screening approach on the mechanisms and consequences underlying APA regulation (12). The deployment of such a portal can provide immediate usability of the resources showcased in publications, and thus increase their impact by releasing a more user-friendly version of the data, thus reaching a wider number of interested researchers.

The R/Shiny framework is highly flexible in regard to data manipulation and analysis, therefore we expect straightforward extensions and adaptations to the scenario we proposed in this work, also to other experimental settings. As computational reproducibility gains more attention (94; 95; 96) in the scientific community, it will be desirable to support the creation of a report-like document, e.g. implemented with knitr/R Markdown, tracking the performed analyses, similar to the approach followed by the pcaExplorer package (97).

Finally, we anticipate TREND-DB to become a rich resource to explore and validate the impact of genetic and epigenetic variants on APA, and thereby also help to expand our understanding of the genetic origins of disease (98).

## ACKNOWLEDGEMENTS

The authors would like to express their gratitude to current and former members of the Danckwardt lab.

## SUPPLEMENTARY DATA

Supplementary Data (Supplementary Table 1, RNAi targets and effects on alternative polyadenylation (APA)) and Supplementary Methods are available online.

