## Supplementary Table 1 for "TREND-DB – A Transcriptome-wide Atlas of the Dynamic Landscape of Alternative Polyadenylation"

**RNAi targets and effects on alternative polyadenylation (APA).**

Filtered TRENDseq reads represent the reads aligned to true APA isoforms excluding internal priming events (see Materials and Methods; Ogorodnikov et al. Nature Communications 2018)

| Func category | Gene Name | Number of APA-affected genes | Mean distance between regulated APA sites (nt) | Filtered TRENDseq reads | Total number of APA-affected genes per category | Percent per category | Average effect per category (number of APA-affected genes) | Mean distance between regulated APA sites (nt) per category |
|---|---|---|---|---|---|---|---|---|
| **Cleavage and Polyadenylation (CPA)** | CPSF1 | 492 | 922 | 653905 | | | | |
| | CPSF2 | 158 | 867 | 272105 | | | | |
| | CPSF3 | 164 | 820 | 524221 | | | | |
| | CPSF4 | 407 | 959 | 649031 | | | | |
| | FIP1L1 | 399 | 984 | 596315 | | | | |
| | CSTF1 | 402 | 1065 | 981763 | | | | |
| | CSTF2 | 39 | 1141 | 273094 | | | | |
| | CSTF3 | 536 | 1039 | 508429 | | | | |
| | CPSF6 | 1163 | 1573 | 250611 | | | | |
| | CPSF7 | 126 | 757 | 380527 | | | | |
| | NUDT21 | 1320 | 1445 | 1169866 | | | | |
| | PCF11 | 927 | 1573 | 472071 | | | | |
| | CLP1 | 433 | 1183 | 673019 | | | | |
| | CPSF3L | 280 | 911 | 738161 | | | | |
| | SYMPK | 55 | 999 | 201105 | | | | |
| | CSTF2T | 115 | 938 | 587529 | | | | |
| | WDR33 | 244 | 798 | 417290 | | | | |
| | RBBP6 | 103 | 1124 | 243276 | | | | |
| | CPEB1 | 151 | 1002 | 426169 | | | | |
| | PAPOLA | 299 | 1228 | 528244 | | | | |
| | PAPOLG | 123 | 1032 | 513028 | | | | |
| | PAPD4 | 145 | 986 | 233255 | | | | |
| | PAPD5 | 335 | 1190 | 1074623 | | | | |
| | PAPD7 | 40 | 522 | 239409 | | | | |
| | PABPC4 | 19 | 345 | 205648 | | | | |
| | PABPN1 | 93 | 964 | 204929 | | | | |
| | PABPC1 | 54 | 1506 | 398902 | **8622** | **38** | **319** | **1032** |

| | | | | | | | | |
|---|---|---|---|---|---|---|---|---|
| **Transcription** | TBP | 56 | 895 | 353003 | | | | |
| | E2F4 | 84 | 1024 | 493374 | | | | |
| | NRF1 | 165 | 899 | 490875 | | | | |
| | MEF2D | 108 | 780 | 446531 | | | | |
| | YBX1 | 114 | 788 | 370643 | | | | |
| | CTCF | 226 | 742 | 549801 | | | | |
| | TTF2 | 155 | 749 | 677614 | | | | |
| | POLR2A | 98 | 954 | 307657 | | | | |
| | POLR2B | 235 | 1033 | 450259 | | | | |
| | POLR2C | 87 | 951 | 364246 | | | | |
| | POLR2E | 67 | 811 | 233400 | | | | |
| | POLR2J | 164 | 950 | 468082 | | | | |
| | GTF2I | 43 | 1385 | 240473 | | | | |
| | PAF1 | 229 | 1091 | 424176 | | | | |
| | LEO1 | 149 | 964 | 558264 | | | | |
| | CTR9 | 43 | 688 | 195865 | | | | |
| | CDC73 | 69 | 1140 | 264859 | | | | |
| | WDR61 | 147 | 964 | 480531 | | | | |
| | CDK8 | 98 | 778 | 403788 | | | | |
| | CDK9 | 157 | 1057 | 596008 | | | | |
| | CDK7 | 231 | 830 | 674539 | | | | |
| | PIN1 | 157 | 858 | 602239 | | | | |
| | SSU72 | 221 | 992 | 351630 | | | | |
| | CTDP1 | 84 | 827 | 504107 | **3047** | **13** | **127** | **920** |
| **RNA splicing** | SCAF1 | 284 | 974 | 1036661 | | | | |
| | SNRNP70 | 189 | 1444 | 29708 | | | | |
| | SNRPA | 35 | 1295 | 215148 | | | | |
| | SF1 | 111 | 1028 | 380310 | | | | |
| | PTBP1 | 107 | 1196 | 376635 | | | | |
| | U2AF1 | 290 | 1290 | 593729 | | | | |
| | U2AF2 | 153 | 1254 | 353909 | | | | |
| | SF3A1 | 361 | 1181 | 560390 | | | | |
| | SF3B1 | 30 | 1471 | 109284 | | | | |
| | SNRPA1 | 171 | 1318 | 507196 | | | | |

| | | | | | | | | |
|---|---|---|---|---|---|---|---|---|
| | DDX39B | 67 | 955 | 255561 | | | | |
| | DDX23 | 149 | 944 | 461848 | | | | |
| | SRRM1 | 96 | 1015 | 463756 | | | | |
| | SNRNP48 | 133 | 912 | 551099 | | | | |
| | SRSF1 | 73 | 1108 | 236334 | | | | |
| | HNRNPH1 | 62 | 1050 | 343084 | | | | |
| | HNRNPH2 | 190 | 1219 | 820272 | | | | |
| | HNRNPH3 | 19 | 951 | 247690 | | | | |
| | NONO | 135 | 1073 | 825226 | | | | |
| | SFPQ | 70 | 794 | 183291 | | | | |
| | SRSF3 | 354 | 1277 | 505983 | | | | |
| | SRSF4 | 100 | 1048 | 475615 | | | | |
| | NOVA2 | 32 | 1034 | 306753 | | | | |
| | RBM5 | 46 | 669 | 441644 | | | | |
| | HNRNPA2B1 | 128 | 1159 | 353433 | | | | |
| | HNRNPF | 155 | 1109 | 912358 | | | | |
| | SRRM2 | 68 | 1071 | 418622 | | | | |
| | ZCCHC8 | 74 | 1097 | 535354 | | | | |
| | CDC40 | 25 | 835 | 188326 | **3682** | **16** | **127** | **1105** |
| **RNA turnover** | XRN1 | 55 | 1256 | 612368 | | | | |
| | XRN2 | 147 | 795 | 530047 | | | | |
| | CNOT7 | 71 | 575 | 547525 | | | | |
| | CNOT2 | 111 | 953 | 569155 | | | | |
| | TNRC6A | 89 | 1112 | 389649 | | | | |
| | ZFP36 | 34 | 886 | 365198 | | | | |
| | TTC37 | 72 | 1084 | 326987 | | | | |
| | UPF1 | 32 | 1630 | 360228 | | | | |
| | UPF2 | 37 | 1298 | 396584 | | | | |
| | UPF3A | 66 | 1015 | 416470 | | | | |
| | UPF3B | 131 | 819 | 827286 | | | | |
| | RNPS1 | 269 | 1086 | 790214 | | | | |
| | ALYREF | 214 | 1801 | 509470 | | | | |
| | SMG6 | 103 | 1002 | 467809 | | | | |
| | CASC3 | 35 | 832 | 239502 | | | | |

| | | | | | | | | |
|---|---|---|---|---|---|---|---|---|
| | DIS3 | 42 | 1113 | 369092 | | | | |
| | DIS3L | 91 | 987 | 636236 | | | | |
| | EXOSC10 | 70 | 1235 | 428481 | | | | |
| | SKIV2L2 | 22 | 687 | 367144 | | | | |
| | SKIV2L | 96 | 1014 | 611966 | | | | |
| | EXOSC6 | 45 | 1078 | 258519 | | | | |
| | DCP1A | 82 | 925 | 567538 | | | | |
| | DCP2 | 144 | 1248 | 613251 | **2058** | **9** | **89** | **1062** |
| **Translation** | EIF2A | 28 | 230 | 234306 | | | | |
| | EIF3C | 18 | 701 | 164052 | | | | |
| | EIF4E | 45 | 1267 | 297083 | | | | |
| | EIF4A1 | 103 | 982 | 605673 | | | | |
| | EIF4G1 | 77 | 841 | 561227 | | | | |
| | GNB2L1 | 57 | 1001 | 380891 | | | | |
| | CIRBP | 90 | 1133 | 565364 | **418** | **2** | **60** | **879** |
| **Cell-cyle & Replication** | RPA1 | 171 | 1167 | 1084142 | | | | |
| | RPA3 | 34 | 267 | 251700 | | | | |
| | PCNA | 27 | 1381 | 363853 | | | | |
| | CDK2 | 64 | 834 | 368962 | | | | |
| | NRD1 | 137 | 1071 | 423561 | | | | |
| | CCNB1 | 67 | 1273 | 595124 | | | | |
| | BTG2 | 40 | 368 | 324584 | **540** | **2** | **77** | **909** |
| **Epigenetics** | DNMT1 | 45 | 1138 | 294528 | | | | |
| | TRDMT1 | 40 | 938 | 301975 | | | | |
| | DNMT3A | 193 | 1149 | 716870 | | | | |
| | DNMT3B | 84 | 848 | 558425 | | | | |
| | TUT1 | 51 | 903 | 363603 | | | | |
| | PARN | 117 | 1133 | 825793 | | | | |
| | SLBP | 56 | 1291 | 585795 | | | | |
| | SETD1A | 58 | 828 | 556751 | | | | |
| | SETD1B | 34 | 784 | 328492 | | | | |
| | SETD2 | 36 | 843 | 408521 | | | | |
| | KAT2B | 49 | 885 | 340861 | | | | |
| | SMARCA4 | 74 | 1241 | 641204 | | | | |

| | | | | | | | | |
|---|---|---|---|---|---|---|---|---|
| | CHD1 | 50 | 927 | 434190 | | | | |
| | CHD2 | 32 | 802 | 335955 | | | | |
| | RBL2 | 156 | 1076 | 772549 | | | | |
| | SUPT16H | 40 | 1007 | 386456 | **1115** | **5** | **70** | **987** |
| **Genome surveillance** | PRKDC | 46 | 1220 | 453368 | | | | |
| | XRCC5 | 27 | 954 | 396675 | | | | |
| | XRCC6 | 74 | 900 | 588852 | | | | |
| | TERT | 17 | 1888 | 270488 | **164** | **1** | **33** | **1241** |
| **Cancer associated** | BARD1 | 332 | 1059 | 898038 | | | | |
| | SND1 | 42 | 1247 | 331728 | | | | |
| | MYC | 75 | 1053 | 335105 | | | | |
| | TP53 | 95 | 1024 | 527318 | | | | |
| | CDKN2A | 30 | 1084 | 272627 | | | | |
| | CDKN2D | 20 | 799 | 341439 | | | | |
| | DDX20 | 41 | 757 | 479332 | | | | |
| | DDX42 | 66 | 1266 | 265805 | | | | |
| | XPO4 | 44 | 1242 | 423284 | **745** | **3** | **83** | **1059** |
| **Apoptosis associated** | BCL2 | 49 | 1244 | 459697 | | | | |
| | CASP3 | 57 | 1125 | 434096 | | | | |
| | ELAVL1 | 31 | 1659 | 183432 | | | | |
| | TIA1 | 27 | 1210 | 286873 | | | | |
| | TIAL1 | 99 | 1042 | 653513 | **263** | **1** | **53** | **1256** |
| **Others** | KHSRP | 169 | 1059 | 496587 | | | | |
| | TARBP2 | 103 | 963 | 736887 | | | | |
| | IGF2BP1 | 76 | 831 | 393337 | | | | |
| | NCBP1 | 75 | 1076 | 596502 | | | | |
| | AGO2 | 98 | 790 | 502644 | | | | |
| | DICER1 | 129 | 902 | 679683 | | | | |
| | AHCYL1 | 295 | 921 | 594157 | | | | |
| | MORF4 | 167 | 817 | 582827 | | | | |
| | PPP1CA | 45 | 468 | 424662 | | | | |
| | PPP1CB | 19 | 465 | 248714 | | | | |
| | PPP1R10 | 77 | 1627 | 307421 | | | | |
| | MAPK1 | 61 | 1045 | 544299 | | | | |

| | | | | | | | | |
|---|---|---|---|---|---|---|---|---|
| | MAPK3 | 19 | 832 | 229517 | | | | |
| | MAPK8 | 121 | 1277 | 638029 | | | | |
| | MAPK9 | 18 | 440 | 299948 | | | | |
| | MAPK14 | 15 | 768 | 218768 | | | | |
| | SETX | 98 | 1146 | 392981 | | | | |
| | JUP | 50 | 1610 | 373126 | | | | |
| | RBM7 | 43 | 1313 | 363586 | | | | |
| | IFIT1 | 48 | 966 | 412018 | | | | |
| | IFIT3 | 58 | 847 | 587306 | | | | |
| | FUBP3 | 29 | 1166 | 300709 | | | | |
| | NUSA | 12 | 1032 | 274286 | **1825** | **8** | **79** | **972** |
| **median** | | **130** | **1021** | | | | | |
| **sum** | | **22644** | **100%** | | | | | |
| CPA only | | 8622 | 38% | | | | | |
| all without CPA | | 14022 | 62% | | | | | |
