## Supplementary Methods for "TREND-DB – A Transcriptome-wide Atlas of the Dynamic Landscape of Alternative Polyadenylation"

**Supplementary Methods - Marini F. et al. NAR 2020**

To monitor poly(A) site changes at bona fide functional polyadenylation sites, we first generated a TREND annotation assembly (PMID: 30552333). To this end we employed a tailored approach based on RNA 3′ region extraction and deep sequencing (3'READS), a method which addresses the internal priming and oligo(A) tail issues that commonly plague poly(A) site identification (PMID: 23241633). Importantly this is mainly achieved through an experimental sequence of biochemical wet-lab steps. Briefly, total RNA was subjected to one round of poly(A) selection using oligo(dT) beads, followed by fragmentation on-bead with RNase III. Poly(A)-containing RNA fragments were isolated using streptavidin beads coated with a 5′ biotinylated chimeric dT45 U5 oligo, followed by washing and elution through digestion of the poly(A) tail with RNase H. The part of the poly(A)-tail annealed to the U residues of the oligo is refractory to digestion and thus served as "proxy/surrogate" for the poly(A) tail. Eluted RNA fragments were purified by phenol-chloroform extraction and ethanol precipitation, followed by sequential ligation to a 5′-adenylated 3′-adapter, and subsequent deep sequencing. The resulting reads were aligned to the genome (hg38), and those with at least two non-genomic adenosines at the 3′ end were considered as PolyA Site Supporting (PASS) reads. These were used for the poly(A) site analysis (below).

In a second step, this annotation of bona fide functional poly(A) sites was used to curate the sequencing data obtained with TRENDseq that allowed for massively parallel sequencing in the siRNA screening. Here, mapped reads were filtered from internal priming on genome-encoded adenosine rich regions using the assembly of TREND annotation (see above). The number of reads associated with each TREND isoform was calculated using HTSeq-count, and served as a proxy for the expression of an individual 3′end isoform (and measure for APA regulation accordingly).
